# UC Irvine’s Brain Initiative Cell Atlas Network (BICAN) Brain Procurement Program for the Center for Multiomic Human Brain Cell Atlas Project

**DOI:** 10.64898/2025.12.23.696304

**Authors:** Jillian V. Berry, Bryan M. Tran, Ginny Wu, Zhiqun Tan, Elizabeth Mendoza, Miyoung Kim, Zeynep Arat, Payton Sho Huai Lin, Lorena Valenzuela, Todd C. Holmes, Anna Virovka, Melina Thaxton, Matthew Nguyen, Sabrina Stein, Victoria Raha Sigaroudi, Alexis De La Rosa, Brianna Gawronski, Justine Silva, Lourdes Gonzalez, Sierra Wright, Kevin Wood, Gabriel K. Hui, Darren Anousaya, Vishwaas Reddy Gangeddula, Noah Gama, Jasmine Gama, Thomas Gross, Blynn Bunney, Richard Stein, Pedro Adolfo Sequeira, Firoza Mamdani, Preston Cartagena, William E. Bunney, Jerry Lou, Hannah Lui Park, Amanda Wilson, H. Mark Brooks, Virginia Allhusen, John Crawford, Jason M. Knight, Bing Ren, Joe Ecker, M. Margarita Behrens, C. Dirk Keene, Craig Stark, Elizabeth Head, William Yong, Edwin Monuki, Xiangmin Xu

## Abstract

High-quality neurotypical postmortem human brain tissue is essential but difficult to obtain for constructing comprehensive human brain cell atlases. Here we describe the establishment of UC Irvine’s Brain Procurement Program, a coordinated initiative to collect and process neurotypical donor brains for multiomic mapping studies within the NIH BRAIN Initiative Cell Atlas Network (BICAN) consortium. Through partnerships with the Orange County Coroner’s Office, the UC Irvine Willed Body Program, UCI Medical Center, and the Children’s Hospital of Orange County, we have developed standardized workflows encompassing donor identification, postmortem brain recovery and processing, region-of-interest dissection, neurotypical donor selection, and data management. Our experience demonstrates the feasibility of a community-based, multi-institutional procurement framework while highlighting challenges in recruiting neurotypical donors and ensuring demographic representation reflective of Southern California. We further identify opportunities to strengthen outreach and donation pathways. This program provides a scalable model for advancing population-reflective, high-quality human brain cell atlas efforts.

**Highlights:**

- Establish a multi-site pipeline for procuring high-quality neurotypical human brains.
- Demonstrate feasibility of donor collection across childhood to adulthood.
- Identify barriers and propose strategies to improve broad donor recruitment.

## Introduction

Scientific and medical advances in understanding human brain organization rely on access to high-quality brain tissues ^1^. Despite over a century of progress in human neuropathology and neuroanatomy, the field continues to face persistent barriers in obtaining well-characterized, representative brain samples for human brain studies ^2–6^. Classical brain collections emerged in the late nineteenth century and matured into systematic brain banking programs in the mid-twentieth century, yet the procurement of neurotypical tissue has consistently lagged behind demand. The term “Neurotypical” in our and other studies is used to refers to donors without a documented history of neurological, psychiatric, or neurodevelopmental disorders, and without neuropathological findings inconsistent with typical brain structure or organization for age. High-quality neurotypical human brain tissue remains one of the most limited and scientifically valuable resources in neuroscience. Even today, many U.S. and international brain banks report chronic shortages. These limitations arise not only from the inherently unpredictable nature of postmortem donation, but also from cultural reservations, religious concerns, logistical uncertainties, and mistrust toward medical and research institutions ^4,5,7^. Underrepresented racial and ethnic communities such as African American and Latino groups remain disproportionately excluded due to longstanding inequities in healthcare access, perceived risks, language barriers, and insufficient community outreach ^8,9^. As described before ^10^, recent media reporting on misconduct within parts of the for-profit organ and tissue donation industry has further heightened public caution. These challenges underscore the need for transparent, community-centered pathways for ethically responsible brain donation ^11^.

At the national level, the NIH BRAIN Initiative Cell Atlas Network (BICAN, RRID: SCR_022794) plays a central role in addressing the need for high-quality neurotypical tissue. The BICAN is a coordinated U.S.-wide consortium of researchers, and this organization works to establish a collection of normal postmortem human brains that reflect the demographic populations of the United States. The overarching goal of BICAN is to develop the world’s most comprehensive molecular, cellular, and anatomical reference atlas of the human brain ^12^. Such an atlas will enable deeper insight into human brain development and aging, while providing a foundation to advance our understanding of how brain cell types, circuits, and molecular pathways are altered in different diseases and conditions. High-quality neurotypical postmortem human brain tissue is therefore indispensable for achieving BICAN’s mission to map the cell types and structures that underlie human brain function.

As one of the designated BICAN procurement sites, the University of California, Irvine (UCI) Brain Procurement Program associated with the UCI Center for Neural Circuit Mapping ^13,14^ is tasked with collecting 50 neurotypical human brains from donors aged 2–65 years, spanning three age groups: children, adolescents, and adults. The UCI’s approach leverages a unique network of institutional partnerships, including the Orange County Coroner’s Office, the UCI Willed Body Program, the UCI Medical Center, and the Children’s Hospital of Orange County (CHOC). Importantly, Orange County is among the most ethnically diverse regions in the United States, home to large Hispanic/Latino, Asian American, Pacific Islander, and multiracial populations. Thus our UCI program is uniquely positioned to secure population-representative brain tissue to help generate human brain cell atlases that more accurately reflect the demographic makeup of the U.S. population.

In this manuscript, we detail our end-to-end workflow from donor identification, family consent, and procurement logistics to tissue processing, postmortem MRI, neuropathological assessment, and anatomical dissection of regions of interest (ROIs). We also summarize donor demographics, operational experiences across collection sites, and the logistical, cultural, and institutional challenges encountered during neurotypical brain procurement. In addition, we discuss opportunities to strengthen donor recruitment, enhance donation pathways, and expand community engagement to support large-scale neuroscience initiatives such as BICAN. Through this work, our goal is to provide a scalable, population-reflective procurement model that can be expanded to additional sites and cohorts to advances the development of comprehensive human brain cell atlases.

## Results

### Donor Identification and Eligibility Assessment, and Recruitment Pipelines

The UCI Brain Procurement Program is an integral part of our BICAN Center project, UM1 “The Center for Multiomic Human Brain Cell Atlas” [PIs: Joseph Ecker (contact); Margarita Behrens; Bing Ren; Daofeng Li; Xiangmin Xu]. The team plans to procure 50 donated brains; strict inclusion criteria, neuropathological evaluation, MRI-based quality assessments, and molecular quality control (QC) may reduce the final cohort to 12 donors [ideally two males and two females per age group: childhood (2-10 years old), adolescence (13-20), and adulthood (25-65)].

Donor eligibility was determined through a two-phase inclusion/exclusion process. The initial screening phase evaluated information available at the time of death, including age, suspected cause of death, medical history, time on ventilator, and presence of psychiatric, neurological, or substance-use disorders (Supplemental **Table S1**). Demographic characteristics, infectious disease status, and indicators of tissue quality were also reviewed. These parameters aligned with the BICAN consortium–defined criteria and informed whether a decedent was potentially suitable for brain procurement and further evaluation. Candidate donors passing the initial screen underwent a comprehensive secondary review. This included extraction of full medical records (when available), toxicology assays, serological testing for infectious diseases, and a structured interview with next-of-kin to document demographic information, family medical history, and relevant clinical background. Neuropathological examination and postmortem MRI scans were also performed. Final donor selection was based on the absence of major exclusionary neuropathology, satisfactory tissue integrity, acceptable QC metrics (RNA integrity number and tissue pH), and a demographic profile consistent with project goals.

Donor specimens were sourced from four institutional pipelines, each of which follows a defined set of operational procedures. The Orange County (OC) Coroner’s Office collaboration builds upon an earlier partnership between UCI’s Psychiatric Brain Donation Program and OneLegacy (RRID:SCR_004148), the federally designated organ procurement organization for transplantation in the region. The BICAN partnership with the Coroner’s Office and OneLegacy began in August 2023. A UCI brain donation recruiter stationed at the Coroner’s Office reviews daily decedent lists, which include suspected cause of death, limited medical history, next-of-kin contact information, and notes regarding family interest in tissue donation. OneLegacy maintains priority for organ and tissue recovery for transplantation; cases deemed ineligible for transplantation are subsequently considered for research donation. If the family expresses interest, UCI staff conduct the consent process by telephone using secure electronic documentation. After consent, UCI’s autopsy technician performs the craniotomy and brain retrieval, placing the specimen on ice for immediate transport to the UCI biorepository. Follow-up interviews with next-of-kin are conducted within several weeks to collect demographic and clinical-related details using a structured questionnaire as needed.

The UCI Willed Body Program receives more than 200 whole-body donations annually from individuals aged 18 years and older. Collaboration with the BICAN project began in January 2024. Because participants have already consented to whole-body donation, no additional consent is required for brain procurement. The Program staff screen decedents using BICAN criteria and notify UCI personnel when a donor meets eligibility requirements. After approval, program staff coordinate brain removal, and the specimen is transferred to the biorepository. Next-of-kin interviews are then conducted to obtain demographic and clinical information.

The UC Irvine Medical Center (UCIMC) is the largest public university hospital in Orange County and provides an additional recruitment pathway. Collaboration with UCIMC commenced in January 2025. Daily mortality reports generated through the EPIC electronic medical record system identify potential donors aged 2–65 years. Consistent with the OC Coroner’s Office pipeline, OneLegacy retains priority for organ and tissue procurement. If a decedent meets our inclusion criteria, we contact OneLegacy to determine whether the decedent meets their eligibility requirements. If the decedent is deemed ineligible for OneLegacy, we approach the next-of-kin. Upon family consent and HIPAA authorization, UCI coordinators access medical records for clinical characterization. Brain procurement procedures mirror those used at the OC Coroner’s Office.

The collaboration with the Children’s Hospital of Orange County (CHOC) began in May 2025 to support the BICAN consortium’s interest in generating multiomic atlases of neurotypical pediatric and adolescent brains. Patients in the Pediatric Intensive Care Unit with imminent risk of death are identified by the care team, especially when families have previously expressed interest in autopsy. CHOC’s dedicated staff conduct the parent consent and, when applicable, assent processes. Per CHOC policy, assent is required for patients of aged 7 and older who are deemed to have decision-making capacity. Autopsy personnel perform brain recovery immediately after death, and UCI personnel retrieve the specimen for transport to the biorepository. CHOC staff collect demographic and clinical information using a modified questionnaire, uploading the data directly into REDCap (Research Electronic Data Capture; RRID:SCR_003445), the secure UCI-hosted data management platform.

### Postmortem Brain Collection and Processing

Once a donor was identified as meeting the initial inclusion criteria, the postmortem brain collection workflow (**Figure 1**) was initiated. All procedures were conducted in accordance with ethical and legal requirements. Informed consent including the language permitting unrestricted data sharing for research, was obtained prior to any procedures. Following consent, the recovery team was mobilized and our autopsy technician conducted the brain removal according to our standardized protocol (STAR Methods; see below). Data collection forms documented key variables including time of death and time to autopsy, allowing calculation of the postmortem interval.

**Figure 1.**
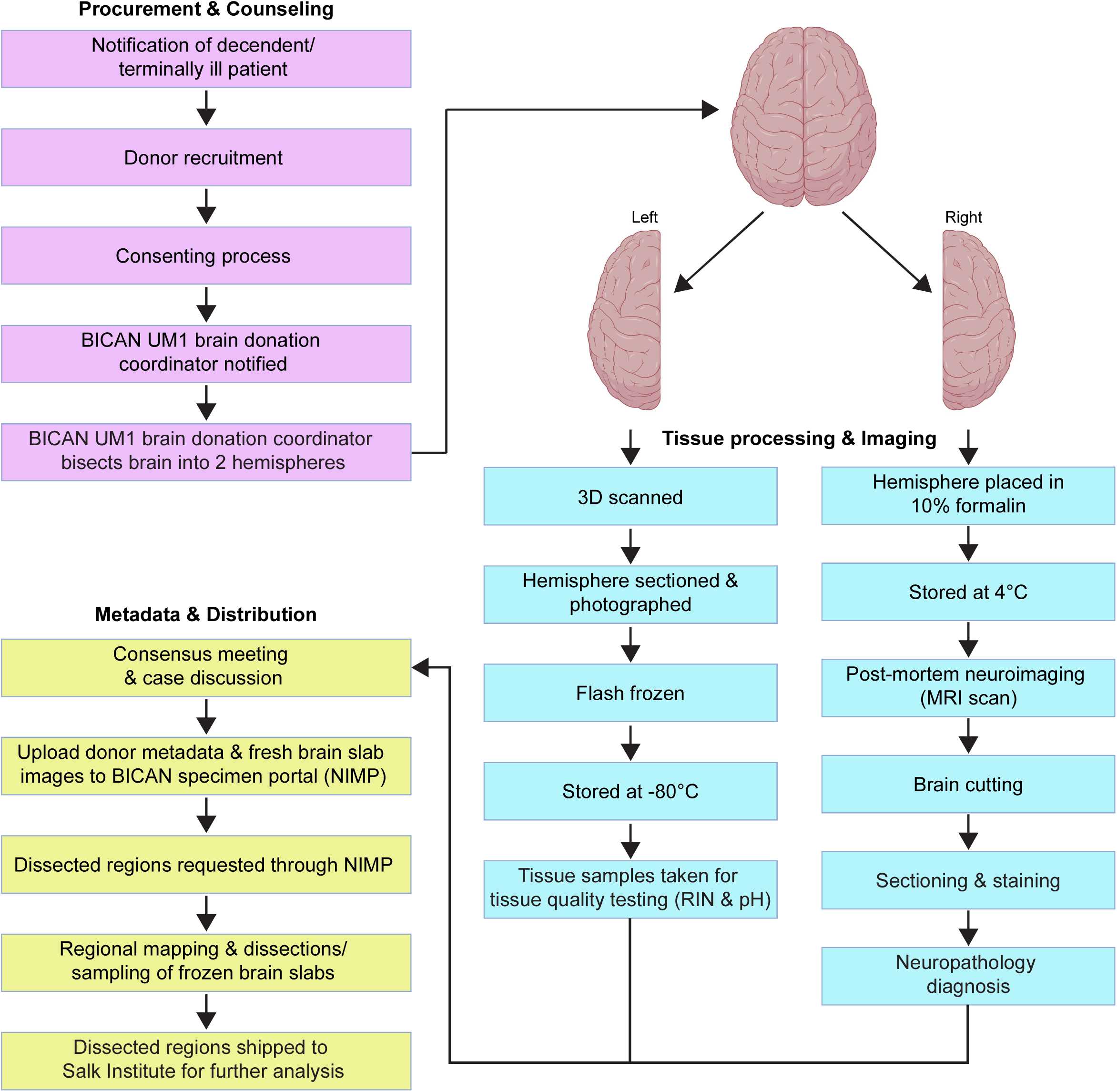
UC Irvine Brain Procurement Program’s pipeline for obtaining brain donations and tissue processing. If a donor meets the initial inclusion criteria, informed consent is obtained and a brain autopsy is performed. The brain is then divided into hemispheres. One hemisphere is sectioned, photographed, flash-frozen, and stored. The other hemisphere is fixed with formalin, imaged using MRI, sectioned, and stained for neuropathological evaluation. When a brain of interest is identified, a consensus meeting is convened to review clinical data, neuropathological findings, and tissue quality. A final determination regarding study inclusion is then made based on established inclusion and exclusion criteria. Metadata and frozen brain slab images from selected neurotypical donors are uploaded to NIMP to be available for request. ROI’s are then mapped through a collaborative effort between UCI and Salk Institute’s BICAN UM1 team. ROI dissections are performed by UCI’s neuropathology and neuroanatomy team, and the resulting tissue is shipped to the SALK Institute for further downstream analysis and cellular/molecular mapping.

Brains procured from all four pipelines were processed according to the identical protocol. After brain removal, the whole brain was weighed, and blood was drawn via cardiac puncture. A portion of the blood sample was submitted to the UCI clinical laboratory for serological testing for hepatitis B virus, hepatitis C virus, and HIV as a safety precaution; donors testing positive for any of these infections were excluded from the study. The brain and blood samples were maintained at 4°C and transported immediately to the core biorepository on the UCI main campus for processing.

At the biorepository, each brain was weighed and bisected into left and right hemispheres (**Figure 1**). The one hemisphere underwent brain surface scanning, sectioning, photography, flash-freezing, and storage. The other hemisphere was fixed in 10% formalin for approximately five weeks, then transferred to PBS + 0.02% sodium azide for long-term storage and future postmortem MRI imaging, as well as neuropathological evaluation.

As for the surface scanning, the fresh hemisphere was digitally scanned using the SHINING 3D EinScan Pro HD system (**Figure 2**). Pre-dissection 3D surface scanning preserves the brain’s native geometry and morphology, providing a real-world reference that facilitates volumetric reconstruction from 2D photographs of brain slabs ^15^. This approach helps to achieve sufficient registration of tissue samples to the Allen Human Brain Atlas ^16^ (Allen Human Reference Atlas, 3D, 2020; RRID:SCR_017764) that serves as a positional common coordinate framework for mapping adult human brain data generated across the BICAN. Following surface scanning, the hemisphere was embedded in alginate to provide structural support during sectioning (**Figure 3**) and coronally sectioned into uniform 4-mm-thick slabs using a custom-designed slicing apparatus ^17,18^. A posterior dot is applied to the medial dorsal surface of each tissue slab using India ink to preserve orientation after freezing (**Figure 3G**). Each section was carefully transferred onto a Teflon plate and digitally photographed to document gross anatomical features and sectioning orientation ^18^. The tissue slabs were then rapidly flash-frozen in a 2-methylbutane and dry ice slurry maintained at approximately -78.5 °C (**Figure 3E-G**) to preserve tissue morphology and molecular integrity for subsequent histological and molecular analyses. Frozen slices were vacuum-sealed in labeled plastic bags and stored at -80 °C (**Figure 3H**). Portions of frozen tissue were sent to the Salk Institute for molecular analyses, including RNA Integrity Number (RIN) assessment ^19^ and genomic sequencing. Additional samples were used at UCI for quality control, including pH testing ^20,21^. The other hemisphere, following fixation in formalin, was subjected to high-resolution MRI prior to systematic sectioning and subsequent anatomical examinations and histological staining for comprehensive neuropathological evaluation (Supplemental **Table S2**). MRI-based neuroimaging has proven to be a highly valuable tool for characterizing brain structure in an intact, non-destructive manner. In particular, multimodal MRI of postmortem human brain tissue ^22,23^ provides a robust approach for assessing global and regional structural integrity, enabling the detection and quantification of structural alterations such as age-related cerebral atrophy and other pathological changes. This hemisphere serves as a long-term reference for structural and histopathological analyses. The remaining tissue from the frozen hemisphere, along with the fixed hemisphere tissue, is stored in the UCI Brain Tissue Repository for additional studies by BICAN investigators and collaborators. Alternatively, in accordance with the requirements of the UCI Willed Body Program, the tissue may be returned to the Program following completion of approved studies.

**Figure 2.**
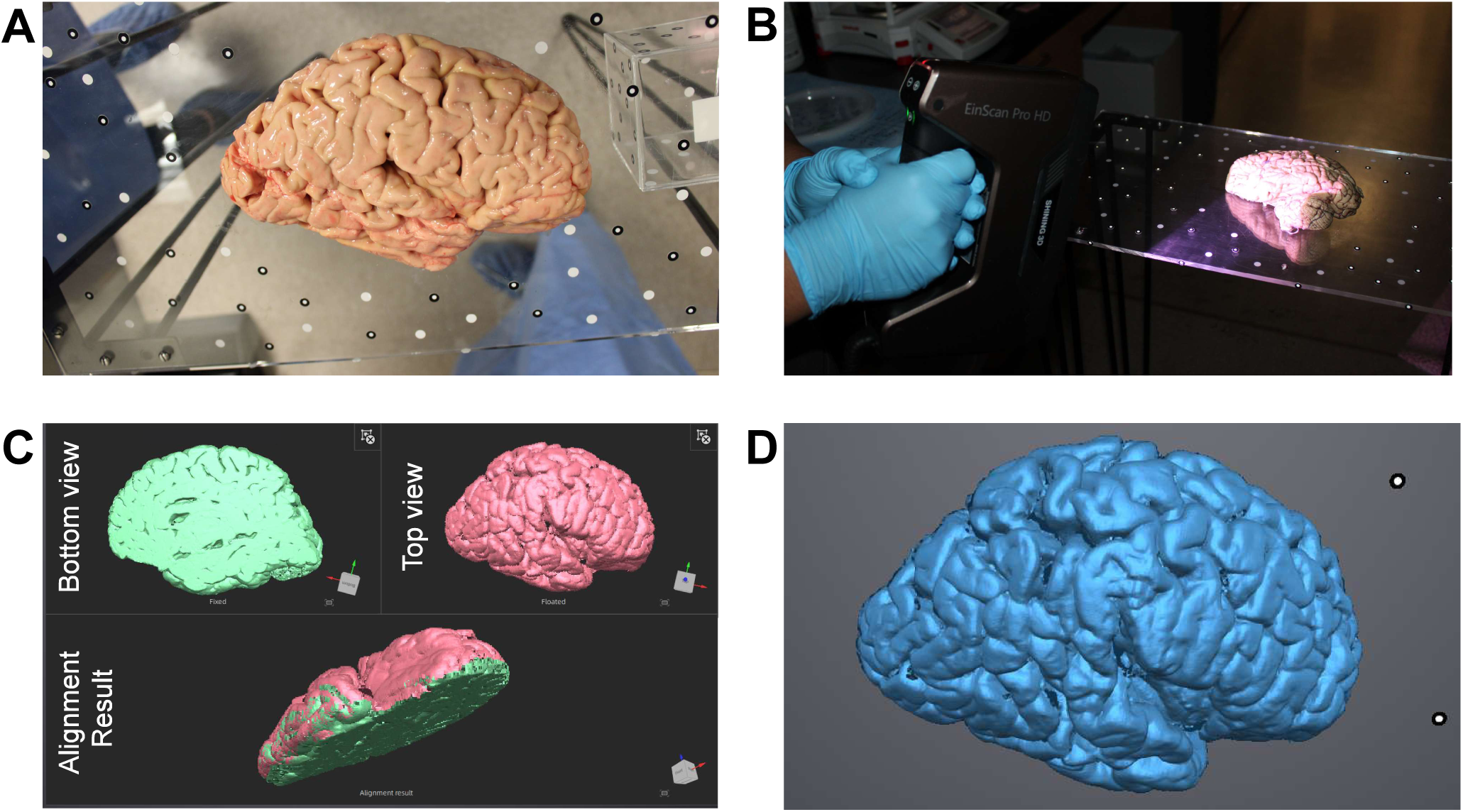
Surface scanning of a human postmortem brain hemisphere. (A) Example of a brain hemisphere positioned on a plexiglass platform for demonstration of the scanning setup. (B) The human brain hemisphere undergoing 3D surface scanning using the CHINING 3D EinScan Pro HD system. (C) Two partial 3D surface scans of a human brain hemisphere acquired by scanning the medial (bottom) face of the hemisphere first and the lateral (top) faces second, with intentional overlap between scans which enable accurate alignment and reconstruction. (D) Final high-resolution 3D surface model generated by merging overlapping scans and applying mesh refinement.

**Figure 3.**
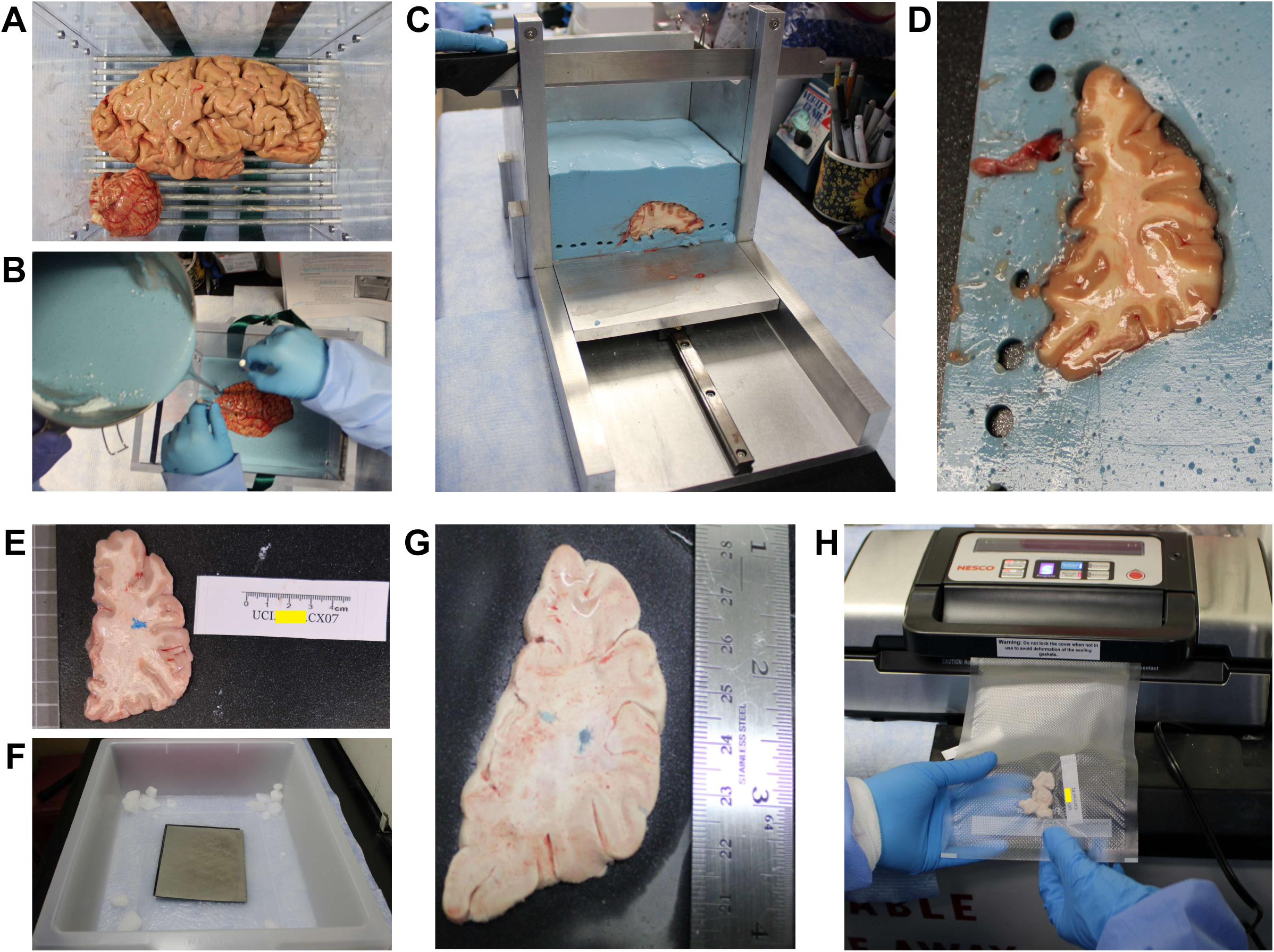
Detailed workflow for brain hemisphere processing. (A) A brain hemisphere (cortex and cerebellum) is aligned in an embedding mold to ensure consistent coronal cuts. (B) Metal rods stabilize the tissue while alginate is poured around it. (C) After solidification, the embedded hemisphere is sliced into 4-mm coronal sections. (D) The sections are transferred to a Teflon plate for individual slabs. (E) An example slab of a hemi-cortex is labeled and marked with a drop of blue dye (India ink) on the posterior surface to indicate orientation. (F-H) The slabs are flash-frozen in a 2-methylbutane and dry ice slurry (F–G), then vacuum-sealed in labeled bags (H) and stored at –80°C for long-term preservation.

### Data Management and Neurotypical Brain Selection

Detailed, de-identified donor information and digital images of brain sections were uploaded to the CUBIE portal built on the Neuroanatomy-anchored Information Management Platform ^24^ (NIMP; RRID: SCR_024684) (**Figure 4**). Developed under the NIH BRAIN Initiative’s BICAN U24MH130988 award as part of the Coordinating Unit for Biostatistics, Informatics, and Engagement (CUBIE), NIMP provides an informatics backbone for collaborative data generation within the BICAN consortium. The Specimen Portal manages tissue resource information of donor/sample registration and images of brain slabs and annotated samples, ensuring direct donor and anatomical linkage of all materials ^24^.

**Figure 4.**
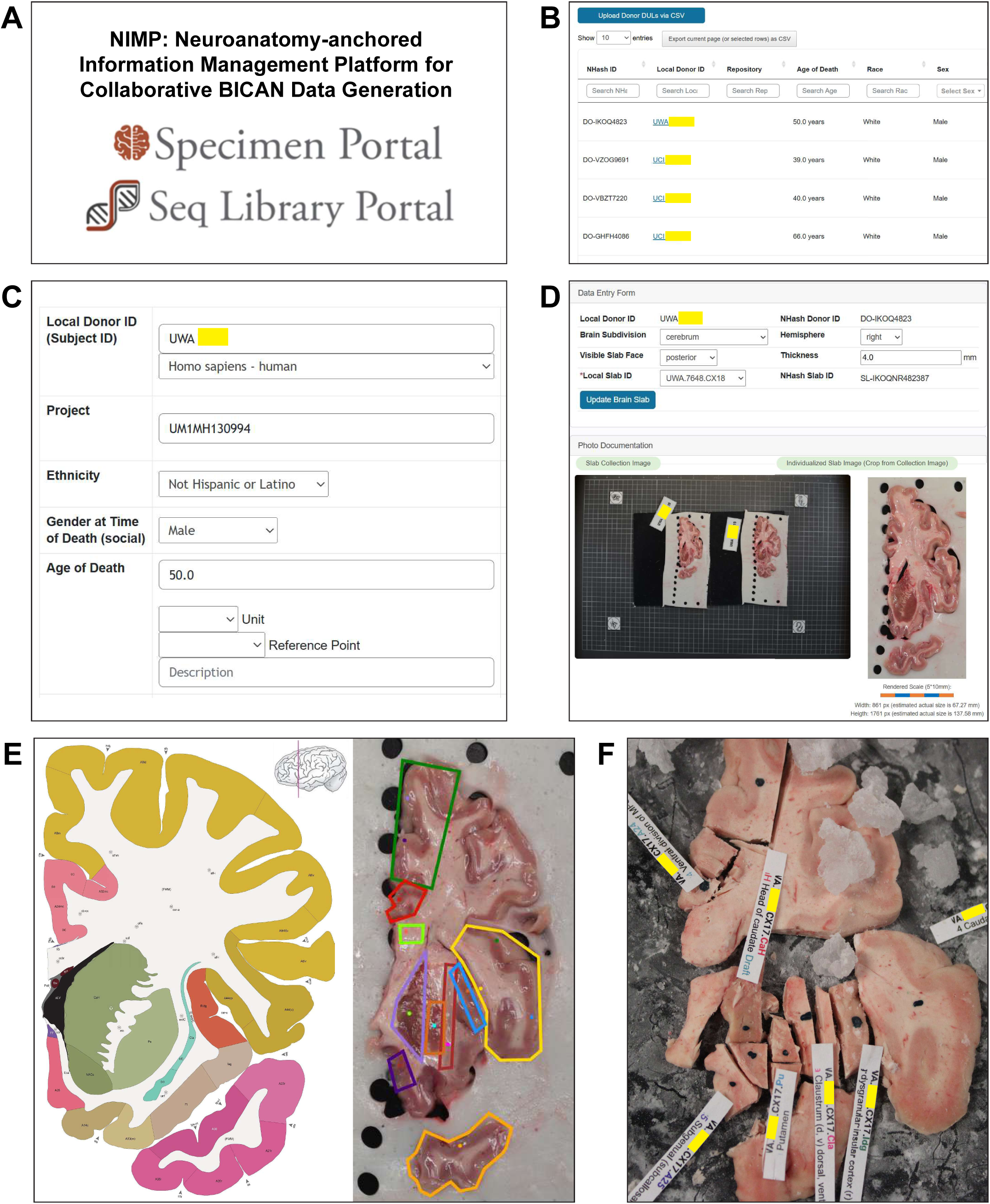
Upload, management, and tracking of donor, brain, and specimen metadata in the NIMP Specimen Portal. (A) Overview of the NIMP BICAN Specimen Portal, including the full program name, acronym, and the two integrated portals: the Specimen Portal and the Sequencing Library Portal and their corresponding logos.(B) Example of a BICAN brain donation team’s or neurobiobank’s donor bank interface which highlights each donor’s clinical, demographic, and specimen metadata and how they are organized within the portal. (C–D) Workflow for uploading donor metadata and brain slab images to the Specimen Portal for review by BICAN researchers and biobank-specific UM1 teams. (E) Portal-based mapping and requesting of regions of interest (ROIs) following donor selection by the research team. (F) The resulting ROI dissections of a specific brain slab based on the integration of approved ROIs into the portal which guide downstream tissue processing.

Once the brain has completed initial processing, the neuropathology and procurement teams convene a formal consensus review to determine eligibility for inclusion in downstream BICAN multiomic analyses ^25^. This evaluation incorporates (1) clinical history and premortem medical records, (2) gross neuropathological inspection (**Figures 2-5**), (3) microscopic evaluation of representative tissue blocks (**Figure 5**), and (4) quantitative quality-control metrics including postmortem interval (PMI), RNA integrity (RIN), pH, and MRI-based structural assessment (**Figure 5**). During the consensus meeting, neuropathological findings are reviewed systematically.

**Figure 5.**
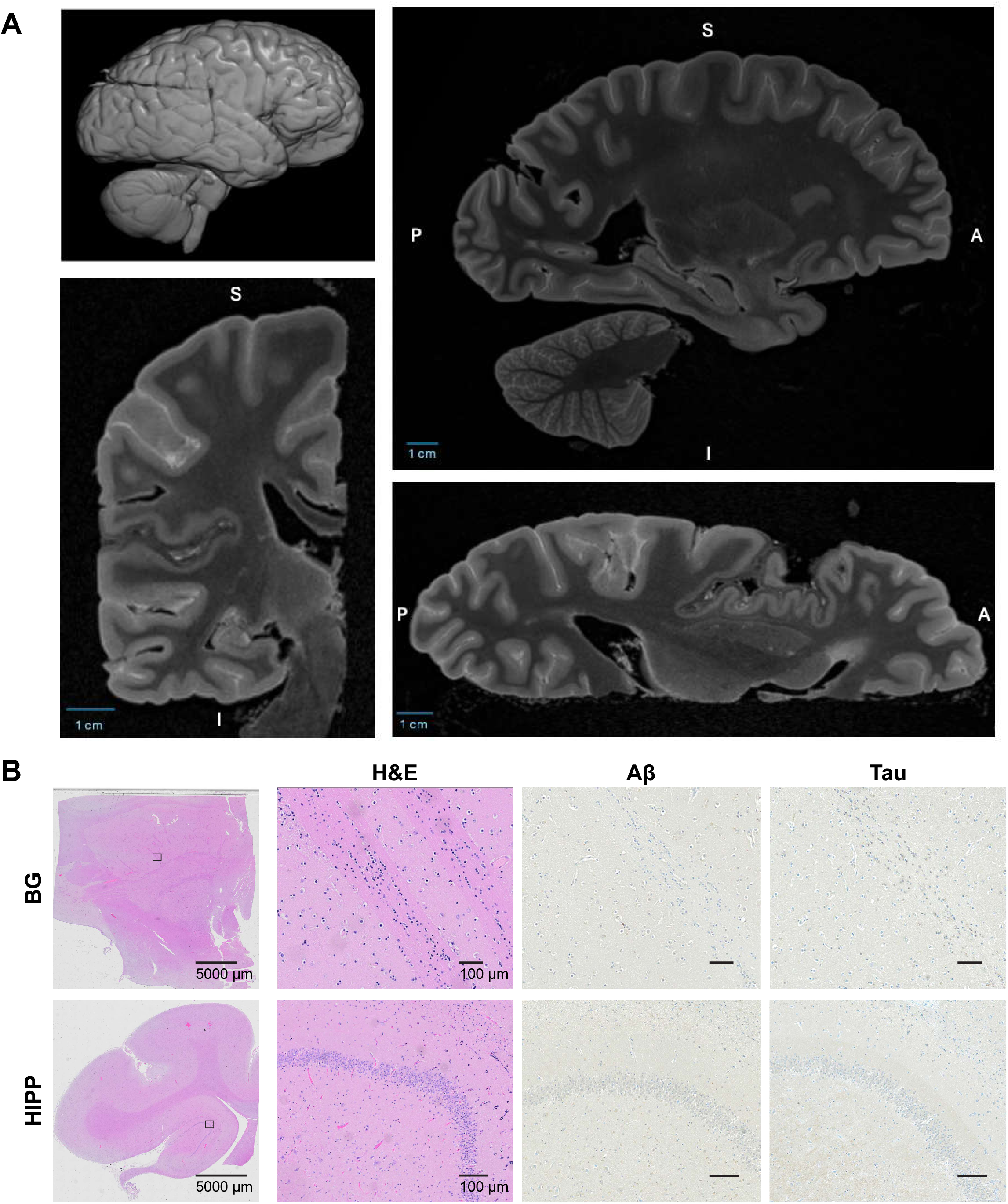
Representative examples illustrating the imaging and tissue-based criteria used in the selection process. (A) MRI datasets acquired during the post-fixation scan are reviewed for structural integrity, distortions, fixation artifacts, or unexpected lesions. (B) Microscopic images of the brain sections stained with hematoxylin and eosin (H&E) stain and selected immunohistochemical markers (Aβ, Tau) are evaluated to assess neurotypical histological characteristics. Basal ganglia: BG; Hippocampus: HIPP.

Representative examples of the data are shown in Figures 2-5, illustrating the imaging and tissue-based criteria used in the selection process. MRI datasets acquired during the post-fixation scan (**Figure 5A**) are reviewed for structural integrity, distortions, fixation artifacts, or unexpected lesions. Gross brain photographs and high-resolution slab images are also examined for evidence of infarcts, contusions, hemorrhage, hydrocephalus, malformations, or other macroscopic abnormalities (**Figures 2-4**). Microscopic sections stained with hematoxylin and eosin (H&E) stain (RRID:SCR_013652) and selected immunohistochemical markers (Aβ (RRID: AB_2565328), human tau/neurofibrillary (RRID: AB_10013724), α-synuclein (RRID: AB_477506), TDP-43 (RRID: AB_615042) R) are systematically evaluated to assess neurodegenerative, inflammatory and vascular pathology (**Figure 5B**). This evaluation is performed for cases involving individuals over 50 years of age and for any cases in which brain pathology is suspected.

The clinical review includes assessment of cause and manner of death, substance exposure, psychiatric or neurological diagnoses, and medical interventions that could impact tissue quality or confound interpretation of neurotypical reference datasets. All data are evaluated against the established BICAN inclusion and exclusion criteria, which require the absence of neurological disease, minimal neuropathological findings, adequate tissue preservation, and demographic suitability for the age-stratified BICAN atlas construction.

Only the brains meeting all clinical, pathological, and quality thresholds advance to comprehensive ROI dissection and downstream single-cell multiomic and spatial transcriptomic analyses. Brains that do not meet criteria due to incidental neuropathology, insufficient quality metrics, or other exclusion factors are documented and excluded from atlas-generation workflows but may be retained for method development and technical training.

### Brain Region of Interest (ROI) Dissection

For single-cell and spatial multiomic analyses, a standardized set of 125 anatomically defined regions of interest (ROIs, see **Supplemental Data, Figure 4E-F**) derived from the Allen Human Reference Atlas, is mapped and dissected from each donor brain through a coordinated workflow between the UC Irvine procurement team and the Salk Institute tissue sample processing team. ROI selection is based on the BICAN consortium’s curated anatomical ontology, which incorporates major cortical areas, subcortical nuclei, limbic structures, and brain stem regions (see **Supplemental Data**).

Prior to ROI dissection, frozen brain slabs stored at –80 °C are transferred to –20 °C for approximately five hours, allowing gradual temperature equilibration. This controlled warming step ensures that tissue remains firm enough for precise microdissection while preventing crystallization artifacts or RNA degradation that may occur with rapid thawing. Slabs are then placed on a chilled dissection platform maintained between –15 °C and –20 °C, using a mixture of wet ice and dry ice.

Neuropathologists and neuroanatomists perform micro-dissections of all mapped regions of interest using atlas-guided maps and slab photographs to maintain consistent anatomical boundaries (**Figure 4 E-F**). ROIs are extracted using sterile, pre-cooled dissection instruments to minimize warming. For each ROI, tissue integrity is visually inspected, and approximate thickness and orientation are recorded. Any deviations from expected morphology such as subtle vascular changes or incidental anomalies are documented for downstream interpretation.

Immediately after microdissection, each ROI is placed into a pre-labeled vacuum-sealable plastic pouch indicating donor ID, ROI code, hemispheric laterality, and dissection date. The samples are vacuum-sealed to prevent condensation and freezer burn, then flash-frozen on dry ice to preserve molecular quality. All ROIs for a given donor are consolidated in dry-ice–filled insulated containers and shipped overnight to the Salk Institute under proper conditions. At the Salk Institute, these ROIs undergo additional quality-control assessments prior to processing for single-cell epigenomics profiling and spatial transcriptomics. The rigor of this ROI dissection pipeline ensures anatomical precision, preserves molecular integrity, and supports scalable integration across the BICAN atlas framework.

### Brain Procurement Outcomes

As of July 31, 2025, the UCI BICAN Brain Procurement Program has procured a total of 32 donors across the four procurement pipelines described above (**Figures 6 and 7, Supplemental Table S3**). For each donor, demographic and mortality information including age, sex, race, ethnicity, and cause of death, was recorded. Of these 32 donors, 27 individuals met the initial inclusion criteria for neurotypical controls. Tissue-quality screening further narrowed this group, with seven donors meeting all standards required for downstream single-cell multiomic analysis. From this qualified group, three adult donors and one adolescent donor were ultimately selected for inclusion in the BICAN Human Cell Atlas project. To complete the initial developmental and adult dataset, one additional adult donor and one adolescent donor were provided by the University of Washington brain bank, and one childhood donor was contributed by the University of Maryland brain bank.

**Figure 6.**
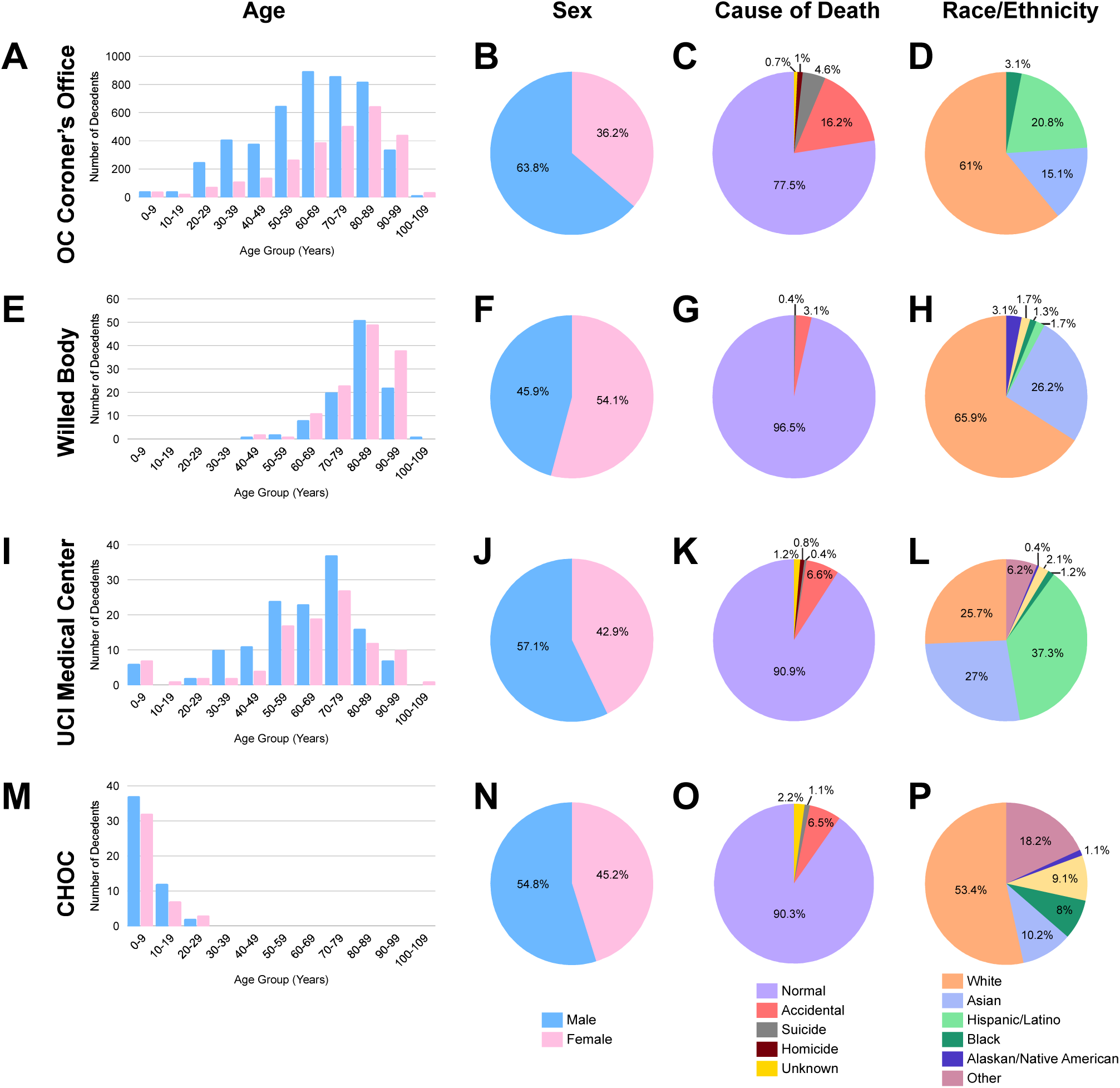
Summary of age and demographic distributions for all decedents screened through each procurement pipeline. (A-D) Orange County Coroner’s Office, (E-H) UCI Willed Body Program, (I-L) UCI Medical Center, and (M-P) Children’s Hospital of Orange County (CHOC). Variables include age at death, sex, race, and ethnicity, providing an overview of the broader populations evaluated for potential inclusion.

**Figure 7.**
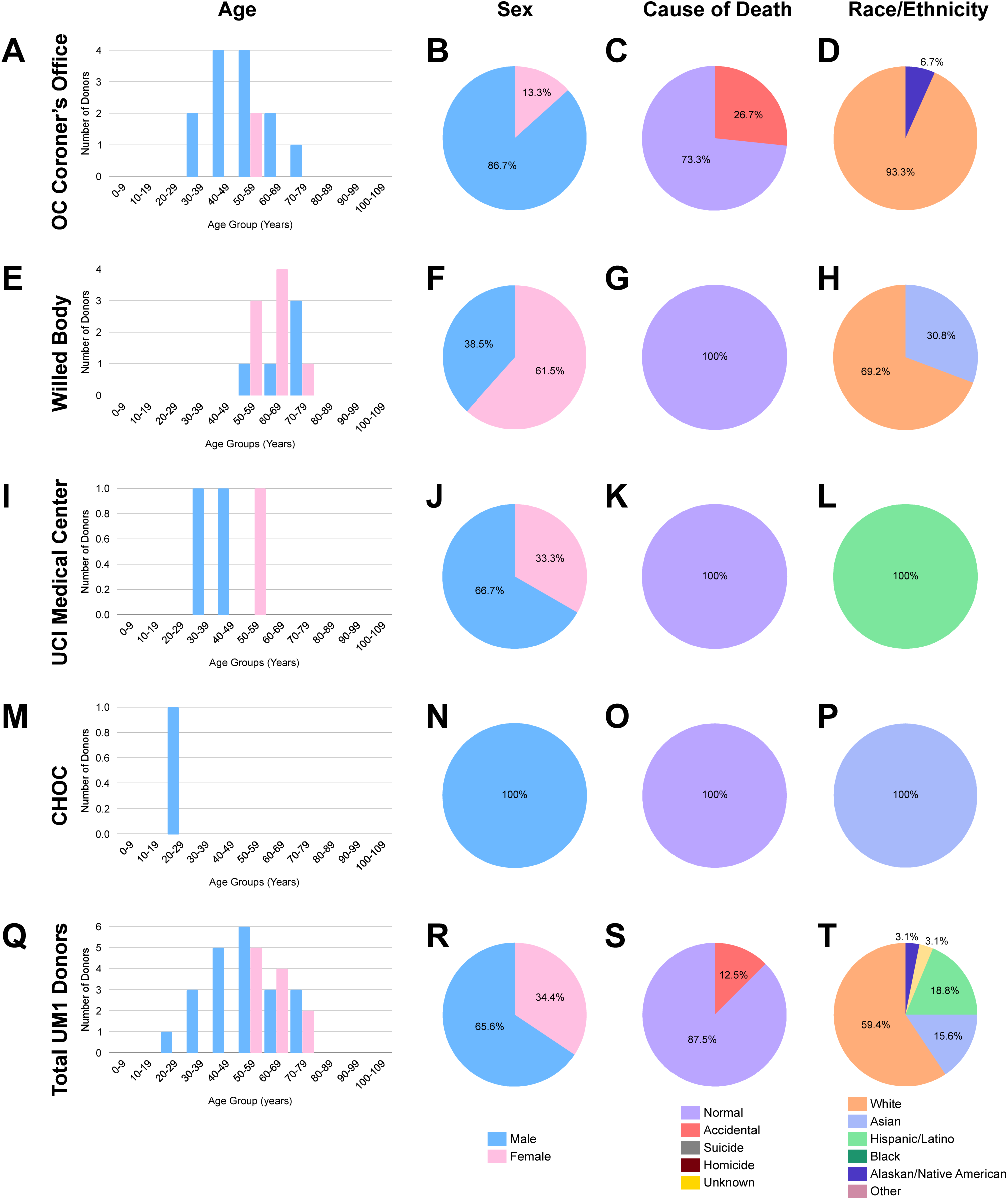
Age and demographic characteristics of BICAN donors obtained from each procurement pipeline. (A-D) Orange County Coroner’s Office, (E-H) UCI Willed Body Program, (I-L) UCI Medical Center, and (M-P) Children’s Hospital of Orange County (CHOC), and (Q-T) total UM1 donors. These data highlight the complementary contributions of multiple pipelines to achieving age and demographic diversity required by the BICAN UM1 project.

The overall demographic characteristics of our donor sources and the 32 acquired donors are summarized in **Figures 6 and 7**. These data illustrate broad variability in sex, race, ethnicity, age, and cause of death across the cohort, reflecting the diversity of the Southern California population and the differing demographic profiles associated with each acquisition pipeline.

### Brain Donor Demographic Information

Demographic and mortality information was systematically collected for all decedents encountered across the four pipelines, including individuals who were screened and those who qualified as BICAN donors. Collected variables included age, race, ethnicity, sex, and cause of death, providing essential context for characterizing both the broader populations evaluated and the subset of donors who met study inclusion criteria (**Figures 6 and 7**).

Donors procured through the Willed Body Program were predominantly older women, which aligns with long-term trends in voluntary whole-body donation programs ^26,27^. Most deaths in this group were attributed to natural causes, and a significant number of exclusions were due to documented neurological or neurodegenerative disease, which was expected given the advanced age distribution of this cohort. Detailed age and demographic profiles of the total decedent pool and UM1 donors from this pipeline are presented in **Figures 6E-H and 7E-H**, respectively. In contrast, donors obtained from the Orange County Coroner’s Office and the UCI Medical Center were more often male and exhibited a wider range of ages and causes of death. The Orange County Coroner’s Office contributed a substantial proportion of younger and middle-aged decedents (**Figures 6A-D** and **7A-D**). Causes of death in this setting frequently included accidents, cardiovascular events, and substance-related causes, reflecting the typical medicolegal population served by the Coroner’s Office. The UCI Medical Center pipeline provided a smaller cohort of potential donors (**Figures 6I-L** and **7I-L**), however, eligible donors from this source were typically accompanied by comprehensive medical records, enabling rigorous assessment of neurotypical status.

The Children’s Hospital of Orange County (CHOC) pipeline contributed one donor: a 20-year-old Asian male who died of leukemia. Although only a single donor was obtained through this pipeline, his inclusion provided essential coverage of the young adult age range within the atlas. **Figure 6M-P and Figure 7M-P** summarize the broader demographic characteristics of decedents screened through CHOC, though the available numbers remain small due to the ethical and logistical challenges associated with pediatric and young adult brain donation.

Collectively, these results demonstrate that a multi-site pipeline approach is necessary to obtain the full range of ages and demographic characteristics required for building a comprehensive human brain cell atlas. The diversity of donors obtained through this program has already contributed significantly to the foundational dataset of the BICAN Human Cell Atlas and provides a strong foundation for future expansion.

## DISCUSSION

Overall, the UC Irvine BICAN Brain Procurement Program demonstrates that a coordinated, multi-institutional, community-embedded framework can overcome many longstanding obstacles to obtaining high-quality neurotypical postmortem human brain tissue. By integrating pipelines from the Orange County Coroner’s Office, the UCI Willed Body Program, UCI Medical Center, and the Children’s Hospital of Orange County (CHOC), we established a robust workflow for identifying potential donors, engaging families, recovering and processing tissue, and applying rigorous clinical, neuropathological, imaging, and molecular quality-control criteria. At the same time, our experience highlights persistent challenges in demographic representation, donor recruitment, and tissue quality that must be addressed to realize the full potential of population-reflective human brain cell atlases.

### Demographic Representation Across Pipelines

Per the U.S. Census Bureau QuickFacts (2023), Orange County is one of the most diverse counties in the United States, with a population that is approximately 36.7% White (non-Hispanic), 34.3% Hispanic/Latino, 24.3% Asian, 2.3% Black/African American, and <1% Native American/Pacific Islander. Our four procurement pipelines collectively access decedent populations that broadly mirror this diversity, but the donors who are ultimately enrolled into the BICAN cohort remain less diverse than the communities from which they are drawn. Each pipeline contributes a distinct demographic profile shaped by its institutional role and catchment area. The UCI Medical Center serves a large, socioeconomically and ethnically heterogeneous patient population and exhibits strong diversity among its decedents (37% Hispanic and 27% Asian), with a relatively young mean age (∼47 years) and balanced sex ratio. Although our donor numbers from UCIMC remain modest, all donors obtained to date identify as Hispanic, underscoring its potential as a critical source for expanding racial and ethnic representation and for recruiting younger adult donors. In contrast, the UCI Willed Body Program primarily contributes older donors (mean age ∼82.9 years) ^28^; their donors are predominantly older, White, and of higher socioeconomic status. These individuals typically pre-register as part of end-of-life planning and often learn about donation through educational or professional networks, which may partially explain the overrepresentation of highly educated White donors ^29^. Recent efforts by the Willed Body Program to engage minority communities such as outreach to Korean organizations in Orange County, have increased Asian representation; currently, 29% of Willed Body donors are Asian, the majority of whom identify as Korean. Women account for 57% of Willed Body donors, likely reflecting sex differences in life expectancy and opportunities for end-of-life planning. As a result, this pipeline has been particularly important for improving representation of female and Asian donors and has contributed several of the highest-quality brains included in the BICAN atlas. The Coroner’s Office presents a markedly different profile. Its decedent population is younger, more heavily male, and encompasses a broad spectrum of causes and manners of death including accidental, homicidal, and undetermined cases. With the exception of sex, the racial and ethnic distribution of decedents more closely approximates the overall county population. However, among donors actually enrolled from the Coroner’s Office, sex and race imbalances remain pronounced: most donors are young males, and nearly all identify as Non-Hispanic White or Hispanic White, with very limited representation from Black/African American and other minoritized groups ^30^. CHOC, our newest pipeline, provides access to pediatric and adolescent populations and therefore covers two of the three key developmental stages (childhood and adolescence) targeted by BICAN. Although we have so far obtained only a single donor from CHOC, its overall decedent demographics are promising, with substantial racial and ethnic diversity and a relatively higher proportion of Black/African American individuals compared with other pipelines.

Taken together, no single pipeline can satisfy the demographic and age targets required for a population-reflective human brain cell atlas. However, the complementary strengths of each pipeline with older female and Asian donors from the Willed Body Program, younger and demographically diverse donors from UCIMC and CHOC, and broad county-wide representation from the OC Coroner’s Office, demonstrate that a multi-site approach can, in aggregate, approximate the racial, ethnic, socioeconomic, and age diversity needed for an equitable human brain atlas.

### Donor Versus Decedent Populations and Recruitment Barriers

A key observation from our program is that the demographics of decedents encountered across pipelines differ markedly from the demographics of donors who ultimately consent to brain donation and meet inclusion criteria. This discrepancy highlights the complex interplay of logistical constraints, medical eligibility, tissue quality, and, most importantly, next-of-kin or donor consent. In the OC Coroner’s Office pipeline, for example, the decedent pool is racially and ethnically diverse, yet nearly all enrolled donors are White, and the vast majority are male. Lower consent rates among families from minoritized communities likely reflect multifactorial barriers, including limited awareness of brain donation, language differences, religious or cultural beliefs, prior experiences with healthcare systems, and broader mistrust of research rooted in historical abuses ^31,32^. Similar patterns are seen nationally in brain, organ, and whole-body donation; recent reports indicate that ∼80% of whole-body donors are White/Caucasian, with disproportionately low representation of Black/African American, Hispanic/Latino, Asian, and Indigenous groups^5,32–34^. Evidence from prior studies suggests that clear, culturally sensitive communication about the purpose of brain donation, how samples will be used, and how donors and their families are respected throughout the process can increase willingness to participate, particularly within African American and Hispanic communities ^5,26,27,32,35–37^. Our experience is consistent with these findings. Specifically, when families have more time to discuss the donation process, ask questions, and understand the potential impact of their contribution, they are more likely to consent.

The UCI Medical Center pipeline, though currently small, illustrates how even a limited number of successful donations from a diverse hospital population can substantially advance representation, by contributing multiple Hispanic donors. Yet recruitment from UCIMC has been constrained by operational overlap with OneLegacy and other transplantation programs, which understandably have priority for organ procurement. To address this, we have initiated a partnership with OneLegacy to identify donors whose organs are recovered for transplantation and whose brains can subsequently be donated for research. This approach has the potential to significantly increase access to high-quality tissue while respecting the primacy of life-saving transplants.

The Willed Body Program operates under a different model: donors make ante-mortem decisions to donate their entire body, including the brain, thus eliminating the need for rapid postmortem consent. This structure simplifies logistics and bypasses acute emotional barriers at the time of death, but it also relies heavily on self-selection by individuals who are already familiar with or positively inclined toward research. As a result, Willed Body programs tend to overrepresent individuals with higher educational attainment and socioeconomic status, again highlighting the importance of targeted outreach to broaden participation ^38,39^.

CHOC presents unique challenges and opportunities. Pediatric and young adult brain donation raises additional ethical considerations, and families may require longer decision-making periods, often leading to higher postmortem intervals. Nevertheless, the ability to recruit donors across childhood and adolescence is crucial for building developmental brain atlases. Strengthening our partnership with CHOC and developing tailored, family-centered communication strategies will be essential for success in this age range ^32,40^.

### Brain Tissue Quality Determinants

In addition to demographic representation, tissue quality remains a critical determinant of whether a donor can be included in the BICAN Atlas project. Consistent with previous literature^19^, we initially anticipated that shorter postmortem intervals (PMIs) would be directly associated with higher RNA integrity numbers (RINs) and more favorable pH values. However, our data did not uniformly support a simple inverse relationship between PMI and tissue quality. Brains procured through the OC Coroner’s Office often had longer PMIs but exhibited relatively high average RIN and pH values, whereas Willed Body donors, despite shorter PMIs, sometimes showed lower molecular quality metrics. These observations suggest that ante-mortem factors including the presence and duration of terminal illness, hypoxia, metabolic disturbances, and agonal state, may exert a larger influence on tissue integrity than PMI alone ^41,42^. Donors who die suddenly from accidents or other acute events, as is common in medicolegal cases, may exhibit better preserved brain tissue despite longer delays before recovery, whereas donors with prolonged systemic disease may show molecular degradation even when recovered quickly. This pattern aligns with prior work indicating that cause of death and agonal factors are critical variables in postmortem tissue quality.

For pediatric donors, additional complexities arise. At CHOC, grieving families may require extended time before consenting to donation, and institutional policies often allow prolonged periods (up to several hours) before brain removal. These delays can modestly reduce RIN and pH values but may be necessary to ensure ethical, family-centered care. Balancing tissue quality against compassionate timing will remain an ongoing consideration in pediatric brain banking.

Our findings argue for a more nuanced approach to tissue-quality assessment that explicitly incorporates ante-mortem clinical status and cause of death, rather than relying on PMI as a primary surrogate. In the long term, a larger donor pool would allow stricter pre-selection based on both clinical and molecular criteria. At present, however, the scarcity of neurotypical donors especially from underrepresented groups and younger age ranges, limits how selective we can be without compromising diversity ^43^.

### Implications for BICAN and Future Directions

As part of the BICAN consortium, our overarching goal is to contribute to a human brain cell atlas that integrates molecular, cellular, anatomical, and demographic diversity. Building such an atlas requires not only high-quality tissue but also deliberate efforts to ensure that donors reflect the racial, ethnic and socioeconomic heterogeneity of the broader population. Our experience shows that this may be achievable through sustained engagement with multiple institutional partners, proactive community outreach, and explicit attention to equity at every stage of the procurement process ^44^.

Several avenues are likely to be important going forward. First, expanding collaborations with additional hospitals, brain banks, neuropathology departments, and organ procurement organizations, particularly those serving communities currently underrepresented in research, will increase both the size and diversity of the donor pool ^45^. The emerging partnership with OneLegacy, the largest organ procurement organization in Southern California, is a key step in this direction and may provide a scalable model for integrating transplant and research donation. Second, ante-mortem donor registration and education should be prioritized wherever possible. Decisions made by individuals during life, rather than solely by next-of-kin under conditions of acute grief, are more likely to reflect informed, sustained consent and may facilitate higher participation from communities that currently show low donation rates. Community-based educational initiatives, collaborations with faith-based and cultural organizations, and inclusion of brain donation information in clinical care settings may all help foster such ante-mortem decisions ^46^. Third, continued refinement of procurement and processing protocols including improved dissection and imaging workflows and optimized preservation methods, will further enhance tissue quality and the utility of collected samples.

In summary, the UCI BICAN Brain Procurement Program provides a comprehensive, community-embedded model for acquiring neurotypical human brains that are both scientifically valuable and increasingly representative of the broad populations. Continued investment in broad recruitment, ethical engagement, and technical innovation will be essential to realizing BICAN’s vision of a human brain cell atlas that benefits all communities.

## STAR Methods

| REAGENT or RESOURCE | SOURCE | IDENTIFIER |
| --- | --- | --- |
| **Biological samples** |  |  |
| Neurotypical adult brain tissue | University of California Irvine Center for Multiomic Human Brain Cell Atlas | Case# UCI 15-24 |
| **Antibodies** |  |  |
| Rabbit polyclonal anti-human tau protein (neurofibrillary tangles) | Agilent (Dako) | RRID: AB_10013724 |
| 6E10 anti-amyloid-β | Biolegend | RRID: AB_2565328 |
| Rabbit anti-α-synuclein | Sigma-Aldrich | RRID: AB_477506 |
| Rabbit anti-TDP-43 | Proteintech | RRID: AB_615042 |
| **Chemicals, peptides, and recombinant proteins** |  |  |
| 10% formalin | Fisher Scientific | Cat# SF99-20 |
| PBS, Phosphate Buffered Saline, 1X Solution, pH 7.4 | Fisher Scientific | Cat# BP243820 |
| Fluorinert | 3M | Cat# FC-3283 |
| HE600 Hematoxylin and eosin (H & E) staining system | Vantana | RRID:SCR_013652 |
| Ventana Discovery XT Immunohistochemistry Processor | Roche | RRID:SCR_018643 |
| **Other** |  |  |
| University of California at Irvine FIBRE Core Facility | UCI FIBRE | RRID:SCR_027777 |
| BRAIN Initiative Cell Atlas Network (BICAN) | https://www.portal.brain-bican.org/ | RRID:SCR_022794 |
| REDCap | REDCap, UC Irvine | RRID:SCR_003445 |
| NIMP: Neuroanatomy-anchored Information<br>Management Platform for<br>Collaborative BICAN Data<br>Generation | https://brain-specimenportal.org/ | RRID:SCR_024684 |
| WashU Human Epigenome<br>Browser | Washington University in St.<br>Louis | RRID: SCR_006208 |

### Key resources table

All brain procurement and processing procedures were conducted under approval from the University of California, Irvine (UCI) Institutional Review Board (IRB #1621, Center for Multiomic Human Brain Cell Atlas). These procedures were performed in collaboration with the Orange County (OC) Coroner’s Office, the UCI Willed Body Program, the UCI Medical Center (UCIMC), and the Children’s Hospital of Orange County (CHOC). We obtained the corresponding decedent data from each of our procurement sites from March 2023 to August 2025.

### Brain Tissue Processing

All procured brains are processed using a standardized protocol. Upon arrival at the UCI biorepository, each brain is weighed, photographed, and bisected into hemispheres. One hemisphere is immersion-fixed in 10% formalin for five weeks and then transferred to phosphate-buffered saline (PBS) containing 0.02% sodium azide for storage and subsequent MRI and neuropathological evaluation. The contralateral hemisphere is processed fresh. Prior to freezing, the surface is captured using a high-resolution SHINING 3D EinScan Pro HD scanner.

The hemisphere is then embedded in alginate to stabilize tissue geometry and coronally sectioned into 4 mm slabs. Each slab is photographed, flash frozen in 2-methylbutane chilled with dry ice, vacuum-sealed, and stored at –80°C. All donor metadata, and slab images are uploaded to the Cubie Brain Specimen Portal (NIMP; RRID: SCR_024684), while clinical and demographic information is stored in REDCap (RRID:SCR_003445). 3D scan images are stored on a secured local drive and are available for request. Frozen tissue is sent to the Salk Institute for RNA integrity assessments, sequencing, and other multiomic workflows; tissue pH is measured at UCI.

### Neuropathological Assessment

A comprehensive neuropathological evaluation is performed in every fixed hemisphere. Gross examination includes assessment of gyral and sulcal patterns, presence of infarcts, hemorrhages, traumatic lesions, ventricular enlargement, and cortical or subcortical atrophy. Representative tissue blocks from cortex, hippocampus, basal ganglia, thalamus, cerebellum, and brainstem undergo routine histological processing. Hematoxylin and eosin staining is conducted using Vantana HE600 system (RRID:SCR_013652) for general morphology assessment. Luxol Fast Blue for myelin; and cresyl violet for neuronal density. Congo red or Thioflavin S staining is performed when amyloid pathology is suspected. Additional immunohistochemical assays are performed on the Ventana Discovery XT Immunohistochemistry Processor (RRID:SCR_018643) using specific antibodies for human tau (neurofibrillary tangles) from Agilent (RRID: AB_10013724), amyloid-β (6E10) from Biolegend (RRID: AB_2565328), α-synuclein from Sigma-Aldrich (RRID: AB_477506), and TDP-43 from Poteintech (RRID: AB_615042) are applied as needed to identify neurodegenerative or inflammatory pathology. A multidisciplinary consensus review integrates neuropathological findings with MRI results, toxicology, and molecular QC to determine eligibility for inclusion in the final donor cohort.

### Region of Interest (ROI) Mapping and Microdissection

The BICAN consortium requires multiomic profiling of 125 anatomical regions. UCI and the Salk Institute jointly map these ROIs onto frozen slabs using MRI and neuroanatomical landmarks. Prior to dissection, slabs are transferred from –80°C to –20°C for approximately five hours to achieve an optimal temperature for microdissection. ROIs are dissected by trained neuroanatomists and neuropathologists, vacuum-sealed, and stored on dry ice until shipment to the Salk Institute for downstream processing.

### Postmortem MRI Acquisition

Formalin-fixed hemispheres undergo high-resolution MRI at UCI’s FIBRE Center using a 3T Siemens Prisma scanner with a 64-channel head coil. Specimens are scanned after a minimum of five weeks of fixation followed by two weeks in PBS with sodium azide. Hemispheres are loaded into a custom-built PVC container (“Conbrainer”) designed to stabilize brain tissue and minimize movement during scanning. The container is filled with Fluorinert (3M, FC-3283) and degassed using vacuum and vibration to remove air bubbles. The specimen is then positioned within the head coil and stabilized using foam supports and sandbags.

Briefly, the Conbrainer is constructed to be a versatile holder not only for human brains, but also for phantoms and for multiple small-animal samples. It was constructed from off-the-shelf PVC/nylon or PLA-based 3D printed parts. A 6” Schedule 40 PVC socket cap and short length of 6” PVC pipe formed the main assembly (bonded with Gorilla heavy duty construction adhesive, model #8212302). A top cap was designed and 3D printed in PLA. The outer diameter of the top cap insert wall is sized to be the same as the inner diameter of the PVC pipe. The maximum diameter of the top-cap face sits flush with the outer diameter of the PVC pipe. On the face of the top cap, two holes are included. Nylon hose mounts are secured to facilitate both filling the Conbrainer with liquid and pulling a vacuum during bubble removal. Additional 3D printed parts include several heights of circular grates to help fill any space not occupied by the tissue. Anchor points are included for securing fiberglass mesh filler that helps secure the sample with minimal distortion, despite the buoyancy of the tissue in Fluorinert.

Each brain hemisphere is placed at the bottom of the container and fastened into place using a fiberglass mesh and two buckle clips. The main volume of Fluorinert is then introduced to the PVC container. A circular grate, along with plastic perforated golf balls, are placed on top of the fiberglass-stabilized brain hemisphere to keep it from floating or moving while remaining invisible to MRI. The top-cap is then sealed to the full completed assembly with Parafilm (Amcor) until mechanically secure against the buoyant forces exerted by the enclosed contents. Using the ports on the top of the Conbrainer, further Fluorinert is added to cover the sample and fill the container. To remove air bubbles, the Conbrainer is placed atop a consumer 3D vibration plate (LifePro 3D Vibration Plate Exercise Machine) with an in-house printed holder and towel used to secure the Conbrainer on the vibration plate from slippage. A vacuum (Kozyvacu KZ350005 8CFM two-stage rotary vane pump) is pulled via one of the two ports (attached also to a liquid trap) and the vibration plate runs on its highest setting for 20 minutes. Approximately once a minute, the Conbrainer is reoriented 10-15 degrees off vertical to different orientations to help dislodge any bubbles. The Conbrainer is then placed inside the 64-channel MRI head coil, secured in place by standard foam pads and sandbags and leveled along two axes. The anterior portion of the head coil is then added. Quick localizer and field map scans are performed to identify any significant bubbles that remain. If observed, the Conbrainer is returned to the shaker table. If not, the main scan is started.

The protocol includes low- and high-resolution structural sequences across multiple contrasts, including field maps, susceptibility-weighted imaging (SWI), T2-weighted FLAIR, T1 3D FLAIR with multiple flip angles, turbo spin echo, T2 SPACE, MP2RAGE, proton density imaging, and multi-shell diffusion-weighted imaging. The specific technical parameters include: Field maps (3 mm isotropic), T2 SWI (0.6 mm x 0.6 x 1.2 mm thickness; 0.6 mm isotropic 2 NEX), T2 FLAIR (0.9 x 0.9 x 2 mm), T1 3D FLAIR (0.8 mm isotropic; 12°, 20°, 28°, and 36° flip angles), T2 Turbo Spin Echo (0.6 x 0.6 x 1.5 mm using TE 48ms, 84 ms, & 120 ms; 0.4 x 0.4 x 2.0 mm TE 55 ms), T2 SPACE (0.8 mm isotropic, 2 NEX; 0.6 mm isotropic, 3 NEX; 0.4 mm isotropic, 3.4 NEX), T1 MP2RAGE (0.8 mm isotropic, 2 NEX), Proton density (0.6 mm isotropic), Diffusion (1.5 x 1.5 x 2 mm 12@b0, 64-direction 4@b=50, 4@b=1000, 4@b=2000, 4@b=6000; repeated with reverse phase encoding). The MRI data support anatomical quality control, facilitate ROI localization, and provide multimodal reference data to accompany histological and molecular analyses.

## DATA AVAILABILITY STATEMENT

The original data and figures supporting this article can be made available by the authors in compliance with the protections for donor confidentiality as outlined in Health Insurance Portability and Accountability Act (HIPAA). All donor metadata, 3D brain surface scans, raw and cropped images of donor slabs have been uploaded and are available in the Coordinating Unit for Biostatistics, Informatics, and Engagement (CUBIE) BICAN Specimen Portal, which is built on the Nauroanatomy-anchored information Management Platform (NIMP;RRID: SCR_024684). Additionally, mapped regions of interest can be accessed and viewed on the WashU Human Epigenome Browser (RRID: SCR_006208). Accompanying clinical and demographic data has been stored in RedCap (RRID:SCR_003445).

In accordance with applicable ethical and legal data sharing requirements, specific data is not publicly accessible. Data will be provided to qualified investigators upon requests directed to the lead contact, Dr. Xiangmin Xu.

## AUTHOR CONTRIBUTIONS

Conceived of Study, X.X.; supervised study, X.X., E.H., E.M.; grant support, M.M.B., J.E., B.R., X.X.; brain procurement and processing, J.V.B., B.M.T., E.M., M.K., Z.A., P.S.H.L, L.V., A.V., V.R.S., A.D.L.R., B.G., J.S., L.G., S.W., K.W., G.K.H., D.A., V.R.G., N.G., J.G., A.S., F.M., P.C., H.M.B., A.W., V.A., J.C., J.M.K., C.D.K; sample dissection, J.V.B., B.M.T., E.M., M.K., Z.A., P.S.H.L, L.V., A.D.L.R., D.A., V.R.G., N.G., J.G., A.S., W.H.Y., E.M.; neuropathology, J.L., W.H.Y.; MRI, A.V., T.G., C.S.; figure preparation, J.V.B., B.M.T., E.M., M.K., P.S.H.L, L.V., T.H., V.R.S., A.D.L.R., D.A., V.R.G., H.L.P., X.X., Z.T., G.W.; manuscript writing and editing, J.V.B., B.M.T., E.M., M.K., P.S.H.L, L.V., T.H., V.R.S., A.D.L.R., D.A., V.R.G., H.L.P., Z.T., G.W., M.M.B, B.B., W.B., X.X.

## ACKNOWLEDGEMENTS

The authors would like to thank the UCI Brain Tissue Repository, and all our brain donors/families for contributing to our project. We would also like to thank additional staff at the Coroner’s Office, The Willed Body Program, UCI Medical Center, and Children’s Hospital of Orange County for collaborating with UCI to allow us to procure donors from their facilities. This work was funded by the NIH grant [UM1MH130994 Center for Multiomic Human Brain Cell Atlas]. This publication was supported and coordinated through the Brain Initiative Cell Atlas Network (BICAN, RRID:SCR_022794).

## Supplemental Materials

**Supplemental Table S1**. Initial inclusion/exclusion criteria for donor eligibility

**Supplemental Table S2**. Anatomical examinations and histological staining for comprehensive neuropathological evaluation

**Supplemental Table S3**. Basic tissue quality-related information for the 32 donors obtained from the four procurement pipelines

**Supplemental Data**: The data sheet of 125 anatomically defined regions of interest for the BICAN UM1 project.

