## Supplemental Tables 1-3 for "UC Irvine’s Brain Initiative Cell Atlas Network (BICAN) Brain Procurement Program for the Center for Multiomic Human Brain Cell Atlas Project"

**Supplemental Table S1. Initial inclusion/exclusion criteria for donor eligibility**

| **Inclusion** | **Exclusion** |
|---|---|
| Age: 2 years to 65 years | Brain related death (including trauma and **brain death**) |
| Terminally ill or recently deceased | Mechanical ventilation > 1 week prior to death |
| All races and ethnicities | Neurologic conditions (dementia, stroke, Parkinson's Disease, epilepsy, MS, etc.) |
| PMI < 30 hours | Prolonged mental illnesses that affect brain function (schizophrenia) |
| Accidental and natural deaths (including acute events without TBI or head trauma) | Brain cancers (and metastatic cancers to brain) |
| | Fentanyl use/abuse/dependence |
| | Positive serology for HIV, Hepatitis B, or Hepatitis C and active SARS-CoV-2 infection at time of death |

**Supplemental Table S2. Anatomical examinations and histological staining for comprehensive neuropathological evaluation**

(A) H&E and other staining

| Block | Brain Region stained with H&E or H&E/LFB | Stain if age >50 or H&E suspicion for pathology |
|---|---|---|
| A1 | Superior/middle frontal gyri (BA9, ACA-MCA watershed, STG incised) | Aβ, Tau/pTau, α-synuclein, TDP-43/pTDP-43 |
| A2 | Inferior parietal lobule (BA39) | Aβ, Tau/pTau |
| A3 | Superior and middle temporal gyri (BA22-BA21, STG incised) | Aβ, Tau/pTau |
| A4 | Occipital lobe (BA17, calcarine and pericalcarine cortex) | Aβ, Tau/pTau |
| A5 | Anterior cingulate gyrus (BA24-BA32) | α-synuclein |
| A6 | Hippocampus at the level of the LGN | Tau/pTau, TDP-43/pTDP-43 |
| A7 | Amygdala | α-synuclein, TDP-43/pTDP-43 |
| A8 | Neostriatum at level of anterior commissure (including caudate, internal capsule, putamen, globus pallidus, claustrum, and extreme capsule) | Aβ |
| A9 | Thalamus with subthalamic nucleus | |
| A10 | Midbrain at the level of the red nucleus | Aβ |
| A11 | Pons with basis pontis and locus ceruleus | |
| A12 | Medulla with inferior olivary nucleus | α-synuclein |
| A13 | Cerebellar cortex with deep cerebellar white matter and nuclei | Aβ |
| A14+ | Gross lesions, radiographic lesions, etc. | |

(B) Potential pathological identifications

| **Characteristics** | **Classifications** |
|---|---|
| **Artifacts** | Gross - brain Removal/Dissection (saw marks, tears, etc.)<br>Microscopic - Autolysis |
| **Developmental Neuropathology** | Brain Malformations |
| **Inflammatory Neuropathology** | Neuroinflammation |
| **Infectious Neuropathology** | Infections |
| **Neoplastic Neuropathology** | Intrinsic or Metastasis |
| **Traumatic Brain Injury** | Acute traumatic brain injury<br>Chronic traumatic brain injury<br>Chronic traumatic encephalopathy - low/high<br>Contusions |
| **Vascular Brain Injury** | Acute Infarcts - Macroinfarcts/Microinfarcts<br>Chronic Infarcts - Macroinfarcts/Microinfarcts<br>Acute Hemorrhage<br>Organizing/Remote Hemorrhage |
| **Neurodegenerative Neuropathology** | Alzheimer's Disease Neuropathologic Change<br>Ax - Thal Aβ Phase number<br>Bx - Braak and Braak Stage number for neurofibrillary tangle distribution<br>Cx - CERAD (The Consortium to Establish a Registry for Alzheimer's Disease)<br>Cerebral Amyloid Angiopathy<br>Lewy Body Disease<br>Limbic-predominant age-related TDP-43 encephalopathy neuropathologic change<br>Hippocampal sclerosis |
| **Age Related Neuropathology** | Cerebrovascular Disease – Arteriosclerosis/Atherosclerosis<br>Primary Age-Related Taupathy (PART)<br>Age-Related Tau Astrogliopathy (ARTAG) |

**Supplemental Table S3. Basic tissue quality-related information for the 32 donors obtained from the four procurement pipelines**

| Pipeline | PMI (Hours) | pH | RIN | | | |
|---|---|---|---|---|---|---|
| | | | Frontal | Occipital | Midbrain | Cerebellum |
| **Coroner's Office (n=15)** | 25.7 (SD = 3.5) | 6.69 (SD = 0.3) | 6.8 (SD = 1.1) | 6.55 (SD = 1.3) | 6.6 (SD = 1.3) | 6.92 (SD = 1.2) |
| **Willed Body (n=13)** | 19.8 (SD = 5.5) | 6.42 (SD = 0.3) | 5.5 (SD = 2.4) | 4.8 (SD = 2.5) | 5.6 (SD = 2.3) | 5.2 (SD = 2.3) |
| **UCI Medical Center (n=3)** | 18.9 (SD = 3.1) | 6.22 (SD = 0.1) | 5.0 (SD = 0.25) | 3.4 (SD = 0.2) | 4.3 (SD = 0.1) | 4.4 (SD = 0.05) |
| **CHOC (n=1)** | 12 | 6.4 | 5.8 | 7.2 | 5.8 | 6.3 |
| **Overall Means (n=32)** | 19.1 (SD = 5.4) | 6.43 (SD = 0.3) | 5.8 (SD = 1.8) | 5.5 (SD = 2.1) | 5.6 (SD = 1.9) | 5.7 (SD = 2.0) |
| **Overall Medians (n=32)** | 23.5 | 6.49 | 7.0 | 6.6 | 5.95 | 6.3 |
| **Interquartile Range (n=32)** | 19.5-26.0 | 6.3-6.8 | 5.0-7.4 | 3.9-7.2 | 4.5-7.6 | 4.9-7.8 |
| **Range (n=32)** | 9-34 | 5.9-7.5 | 1.7-8.4 | 1-7.9 | 1.2-8.6 | 1.4-8.2 |
